# Deciphering the Dynamics of Interlocked Feedback Loops in a Model of the Mammalian Circadian Clock

**DOI:** 10.1101/306852

**Authors:** Dorjsuren Battogtokh, John J. Tyson

**Affiliations:** Department of Theoretical Physics, Physics and Technology Institute of the Mongolian Academy of Sciences, Ulaanbaatar 51, Mongolia; Department of Biological Sciences, Virginia Polytechnic Institute and State University, Blacksburg, Virginia 24061, USA

**Keywords:** circadian rhythm, network dynamics, bifurcation analysis

## Abstract

Mathematical models of fundamental biological processes play an important role in consolidating theory and experiments, especially if they are systematically developed, thoroughly characterized, and well tested by experimental data. In this work, we report a detailed bifurcation analysis of a mathematical model of the mammalian circadian clock network developed by Relogio et al. [16], noteworthy for its consistency with available data. Using one- and two-parameter bifurcation diagrams, we explore how oscillations in the model depend on the expression levels of its constituent genes and the activities of their encoded proteins. These bifurcation diagrams allow us to decipher the dynamics of interlocked feedback loops, by parametric variation of genes and proteins in the model. Among other results, we find that REV-ERB, a member of a subfamily of orphan nuclear receptors, plays a critical role in the intertwined dynamics of Relogio’s model. The bifurcation diagrams reported here can be used for predicting how the core-clock network responds—in terms of period, amplitude and phases of oscillations—to different perturbations.

## Introduction

In the last two decades, there has been tremendous progress in understanding how circadian oscillations are generated and how rhythmicity is sustained in the cells and tissues of many species of animals, plants and fungi [1-7]. It has been established that in all these organisms, core clock genes constitute a ‘circadian network’ that generates endogenous oscillations by repression of the transcription of these clock genes by their own proteins, i.e., by delayed negative-feedback in the transcription-translation-repression loop. Mathematically, these robust rhythms arise as limit cycle oscillations at Hopf bifurcation points [8-10]. Different research groups have implemented this general mechanism in their own unique models of circadian clocks [11-14].

In systems biology, mathematical models are of special utility if they explain a vast amount of experimental data and can be investigated by analytic tools [15]. Recently, Relogio et al. developed a mathematical model of the mammalian circadian-clock network [16], expressed as a set of 19 ordinary differential equations (ODE’s), including 76 kinetic parameters.

The mammalian circadian-clock network drives ∼24 h rhythms in the activity of a heterodimeric transcription factor, CLOCK/BMAL, which drives the expression of other clock genes, including *PERIOD* (*PER*), *CRYPTOCHROME* (*CRY*), *REV-ERBα* and *β* (*REV*), and *RORa, b* and *c* (*ROR*). PER/CRY heterodimers bind to and inhibit CLOCK/BMAL, whereas ROR acts as an activator and REV acts as a repressor of transcription of the *BMAL* gene. Different models of the mammalian circadian clock join together these feedback loops in different ways. Some models view the PER/CRY loop as the master oscillator, with the REV and ROR loops providing fine-tuning and robustness to the oscillations [11-13, 17, 18]. In Relogio’s model, on the other hand, the feedback loops are not hierarchical, but rather they orchestrate the rhythm synergistically. Understanding the dynamics of interlocked negative and positive feedback loops, such as the loops present in the mammalian circadian-clock network, is an important and challenging problem [13, 14, 19, 20].

The main goals of this work are:

- To explore the dynamics of a comprehensive model of the circadian rhythm in mammalian cells (Relogio et al. [16]); a model that has been shown already to be in good agreement with a plethora of experimental data.
- To use bifurcation analysis [21] to disentangle the dynamics of the interlocked feedback loops in Relogio’s model.
- To show how the <u>period</u>, <u>phase</u> and <u>amplitude</u> of circadian oscillations depend on key parameters, namely the maximal rates of expression of five key clock genes (*PER*, *CRY*, *REV*, *ROR*, *BMAL*).
- To use two-parameter bifurcation diagrams to study the interactions between changing levels of expression of <u>pairs</u> of clock genes.
- To provide a foundation for future understanding of circadian control networks, as yet unknown or underappreciated players (see, e.g., [22]) are discovered. In addition, our analysis of the dynamics of the circadian network may be a valuable resource for determining optimal rhythmic signals for the daily synchronization of a variety of cellular processes, especially synchronization of the cell division cycle by the circadian clock [23-25].

Because many future applications of molecular systems biology to human physiology and medicine (e.g., the analysis of clinical data) will require comprehensive mathematical models that describe multiple interlocking control networks in the cell [25-28], we suggest that bifurcation analysis, as practiced here, will be a useful tool for assessing the dynamical properties of these networks within the broader physiology of mammalian cells [29, 30].

## Methods

### Relogio et al. Model of the Mammalian Circadian Clock Network

In Figure 1 we show the wiring diagram of the circadian clock network proposed by Relogio et al. for the murine circadian clock in a single cell [16]. The central node of the network is a heterodimeric transcription factor, CLOCK/BMAL, the yellow box in the center of Figure 1. The mRNAs transcribed from the core clock genes (*PER, CRY, BMAL, REV*, and *ROR*) are represented by blue oval boxes in the nucleus. Clock proteins in the cytoplasm are shown as purple boxes. PER and CRY proteins form multimeric complexes and enter into the nucleus, where they interact with CLOCK/BMAL. The PER/CRY complex represses CLOCK/BMAL-activated transcriptions, thereby creating a delayed negative-feedback loop in the transcription-translation process. We will refer to this delayed negative-feedback loop as the ‘PC loop’, and we will refer to the PER/CRY heterodimeric complex hereafter as the ‘PC complex’. The PC complex is degraded at night, releasing its inhibitory effect on CLOCK/BMAL, to allow a fresh restart of the transcription of clock genes [16].

**Figure 1.**
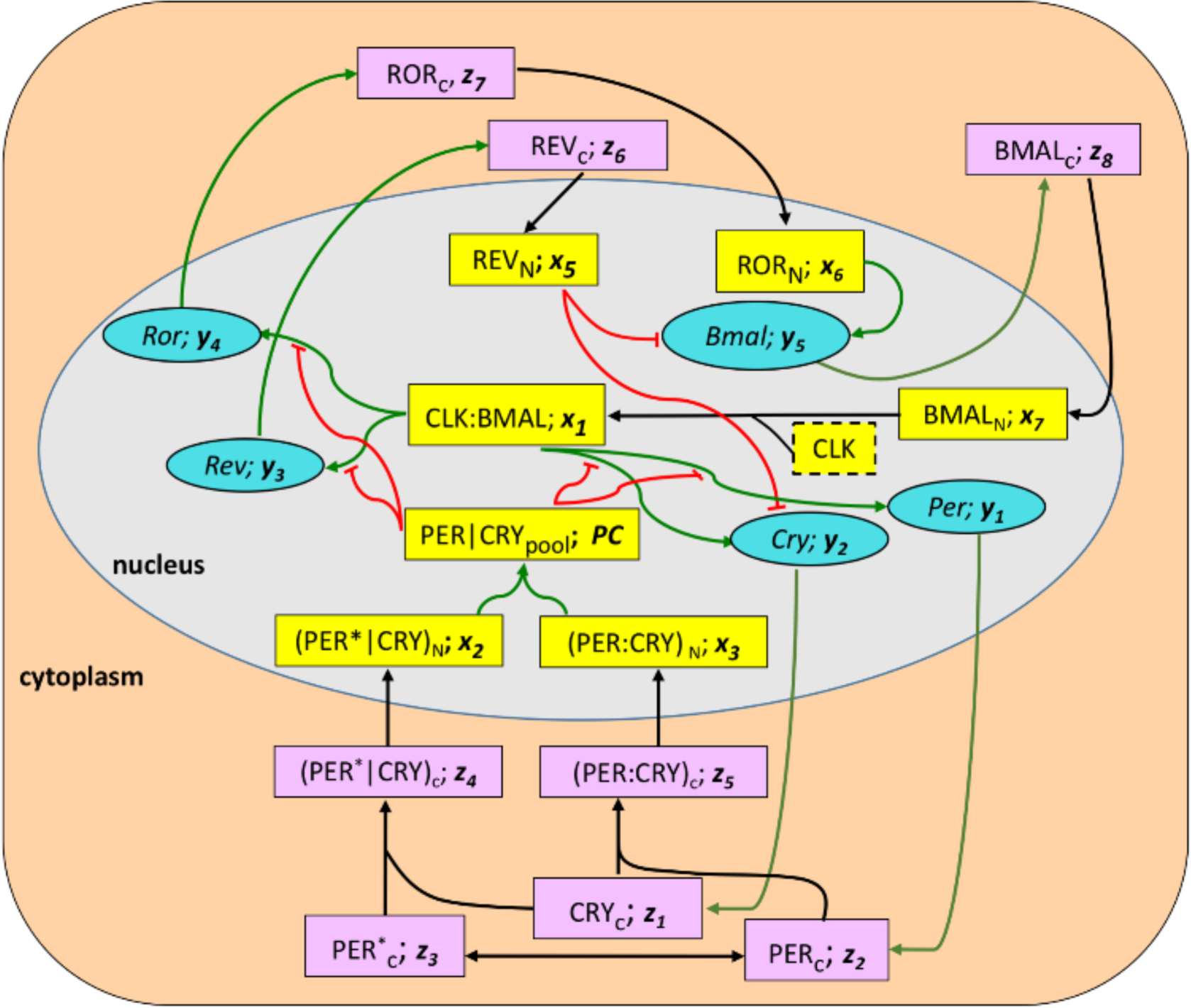
Wiring diagram of Relogio’s model of the circadian clock network in mammalian cells. Core circadian-cloc mRNAs and proteins are indicated by ovals and rectangles, respectively. Within each oval and rectangle is the name of th component followed by the name of its variable in Relogio’s model (Eqs. S1-S19 in Suppl. Text S1). Black arrows indicat transport steps (cytoplasm to nucleus) or chemical reactions (e.g., phosphorylation, indicated by an asterisk*, or associatio of two proteins to form a complex). Green arrows indicate “activation” of one component by another (e.g., the CLK:BMA transcription factor activates synthesis of *Per* mRNA in the nucleus, which in turn activates synthesis of PER protein in th cytoplasm). Red lines, with blunt tips, indicate “inhibition” effects (e.g., nuclear REV inhibits the transcription of *Bmal* an *Cry* mRNAs). In Relogio’s model, CLK protein is assumed to be present in constant excess, so there is no time-dependen variable for this protein.

ROR and REV proteins in the nucleus can bind to the promoter region of the *BMAL* gene and modulate the expression of *Bmal* mRNA. In doing so, ROR acts as an activator, and REV acts as a repressor. Following Relogio et al. [16], we will refer to these feedback effects as the ‘RBR loop’. An important feature of Relogio’s model is that the RBR loops can generate and maintain oscillations autonomously. When the PC loop is maintained constitutively active (non-oscillatory), sustained oscillations continue to be generated by the RBR loops (see the simulations in Fig. 3B of Ref. [16]). Furthermore, when *REV* (or *ROR*) is overexpressed, circadian rhythmicity can be lost (see the simulations in Fig. 3C and Fig. 6A-B of Relogio et al.) [16, 31]. These in *silico* predictions of the model were confirmed by *RORa* and *REV-ERBα* overexpression in a human osteosarcoma cell line, U2OS (see Fig. 7 of Ref. [16]).

**Figure 2.**
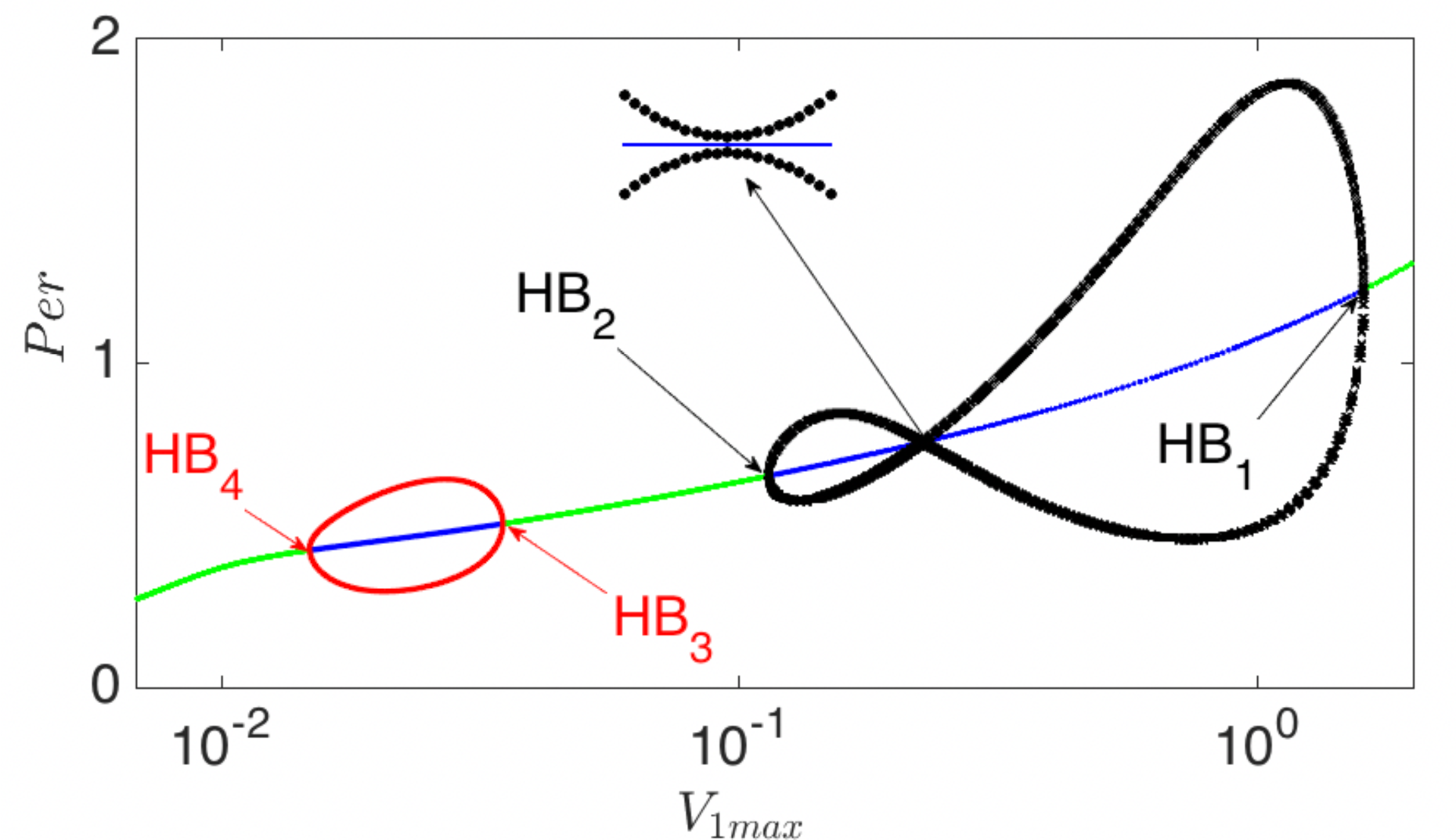
A one-parameter bifurcation diagram of Relogio’s model. The transcription rate of the *PER* gene, *V*_1*max*_, is used as the primary bifurcation parameter. Green lines: stable steady states; blue lines: unstable steady states. There are four Hopf bifurcation points in this diagram, *HB*_*i*_, *i* = 1..4. The black and red curves show maximum and minimum amplitudes of the oscillations.

**Figure 3.**
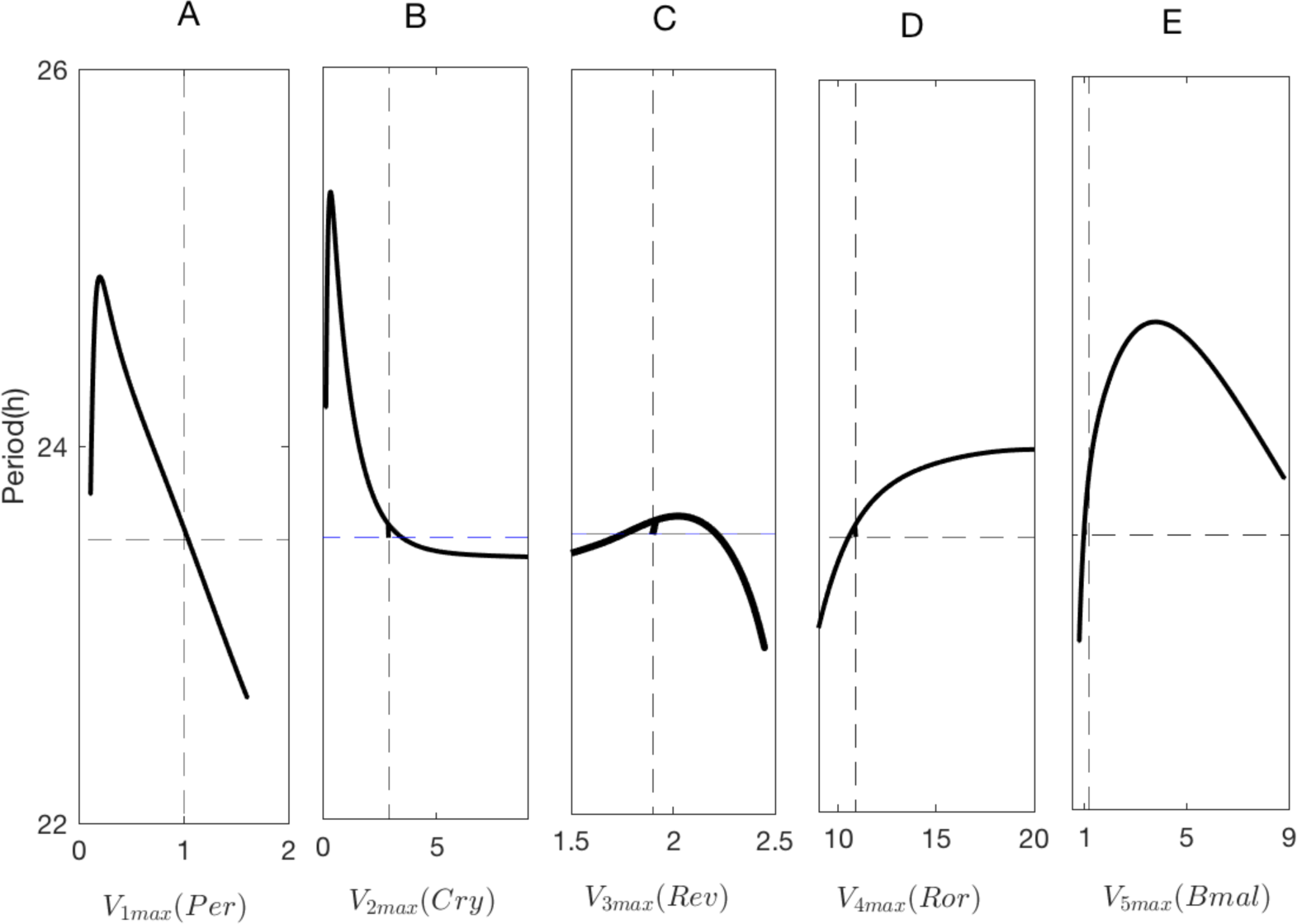
Oscillation period vs *V*_*imax*_, *i* = 1..5, the transcription rates of *Per*, *Cry*, *Rev*, *Ror*, and *Bmal* mRNAs. Th periods of oscillation are plotted between the Hopf bifurcation points *HB*_1_and *HB*_2_in Figure 2. Vertical dashed lines mar the value of the bifurcation parameter corresponding to WT rhythm (period = 23.5 h)

**Figure 4.**
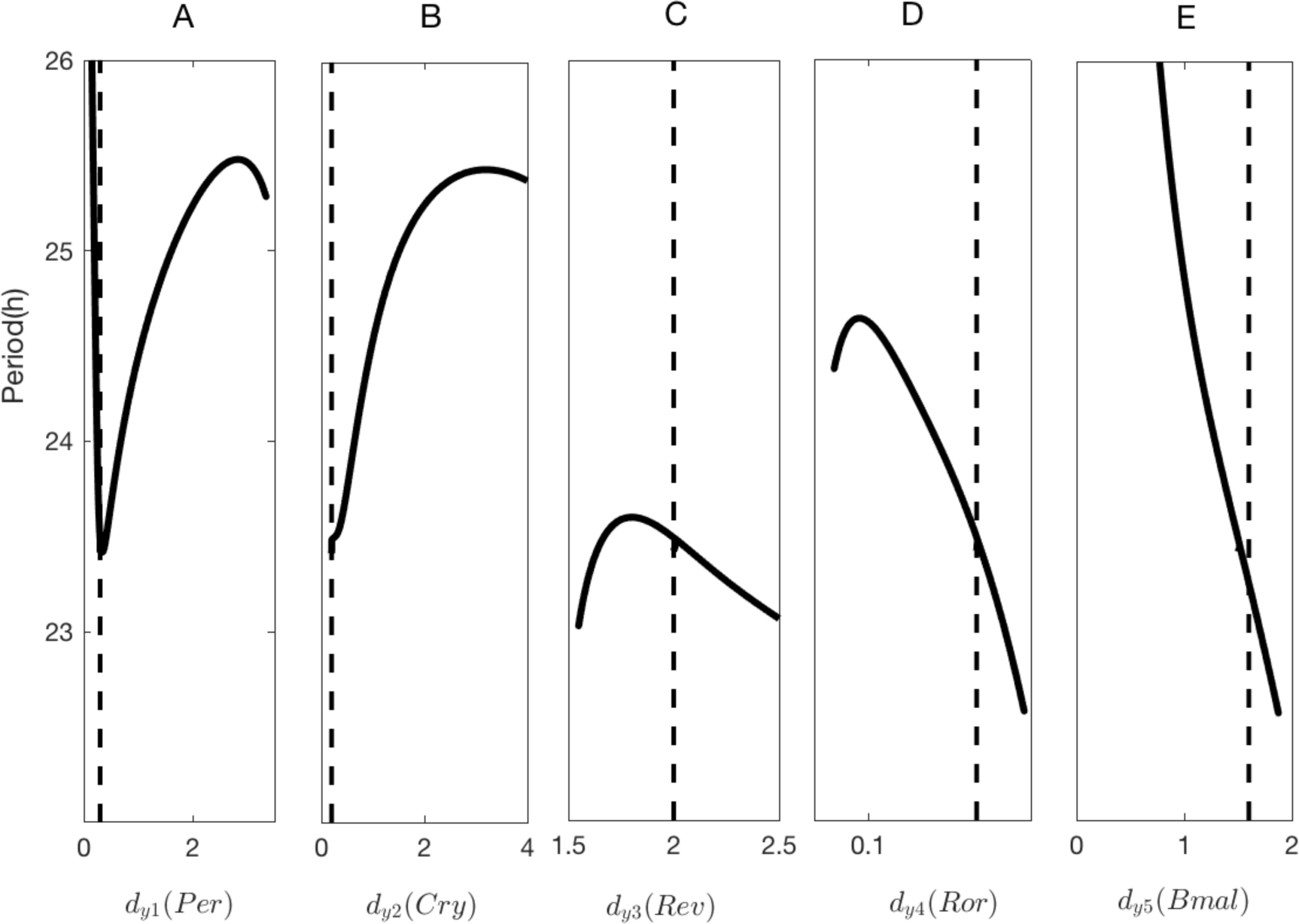
Period vs degradation rates, *d*_*yi*_, *i* = 1..5, of *Per*, *Cry*, *Rev*, *Ror*, and *Bmal* mRNAs. Dashed lines mark WT value of the bifurcation parameters.

**Figure 5.**
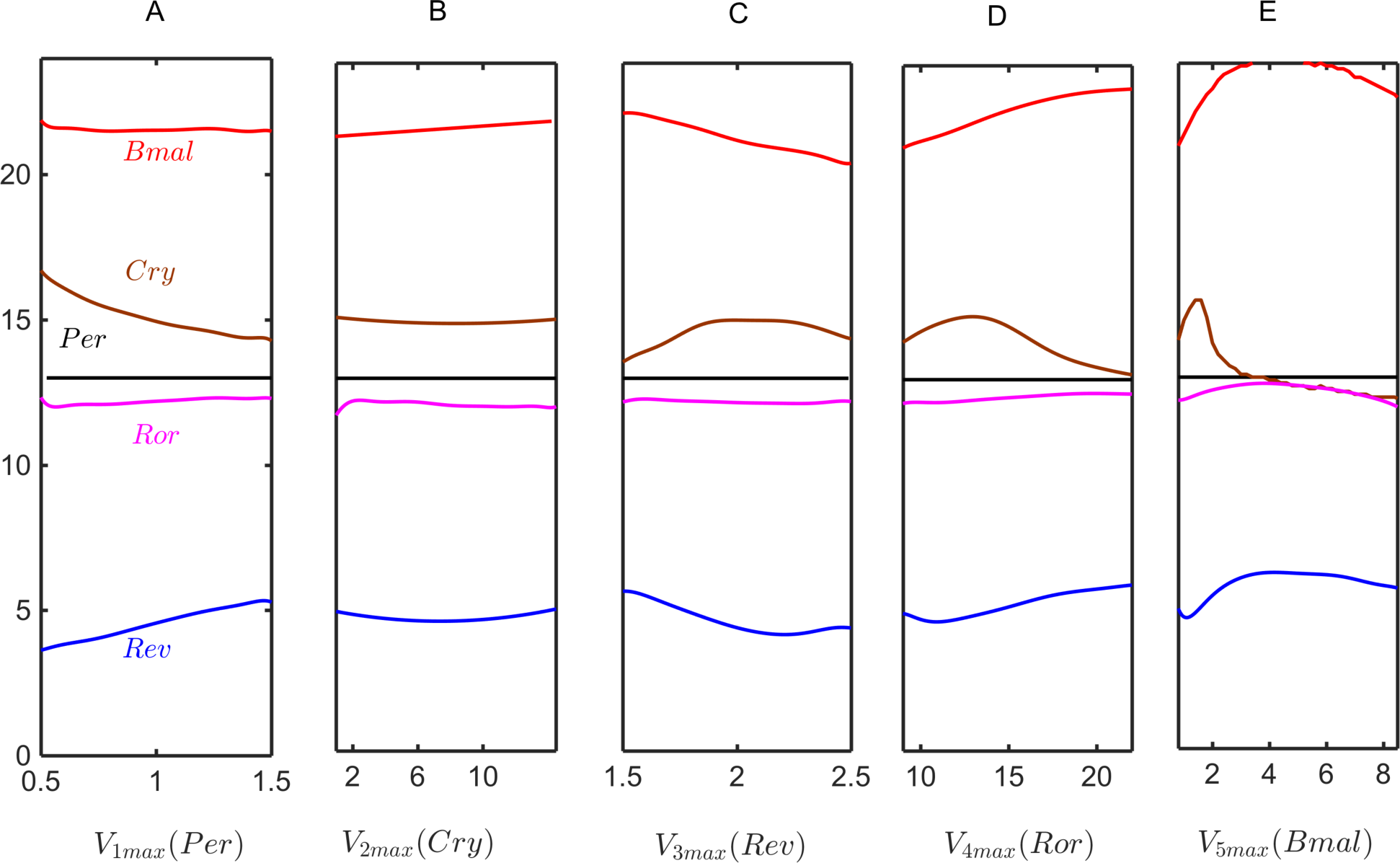
Phases of oscillations (horizontal axis) vs *V*_*imax*_, *i* = 1..5, the transcription rates of *Per*, *Cry*, *Rev*, *Ror*, and *Bmal* mRNAs. Other parameters are fixed at WT values. In these plots, “phase” is the time of maximum expression of the mRNA, relative to the maximum expression of *Per*, which is fixed at 13 h.

**Figure 6.**
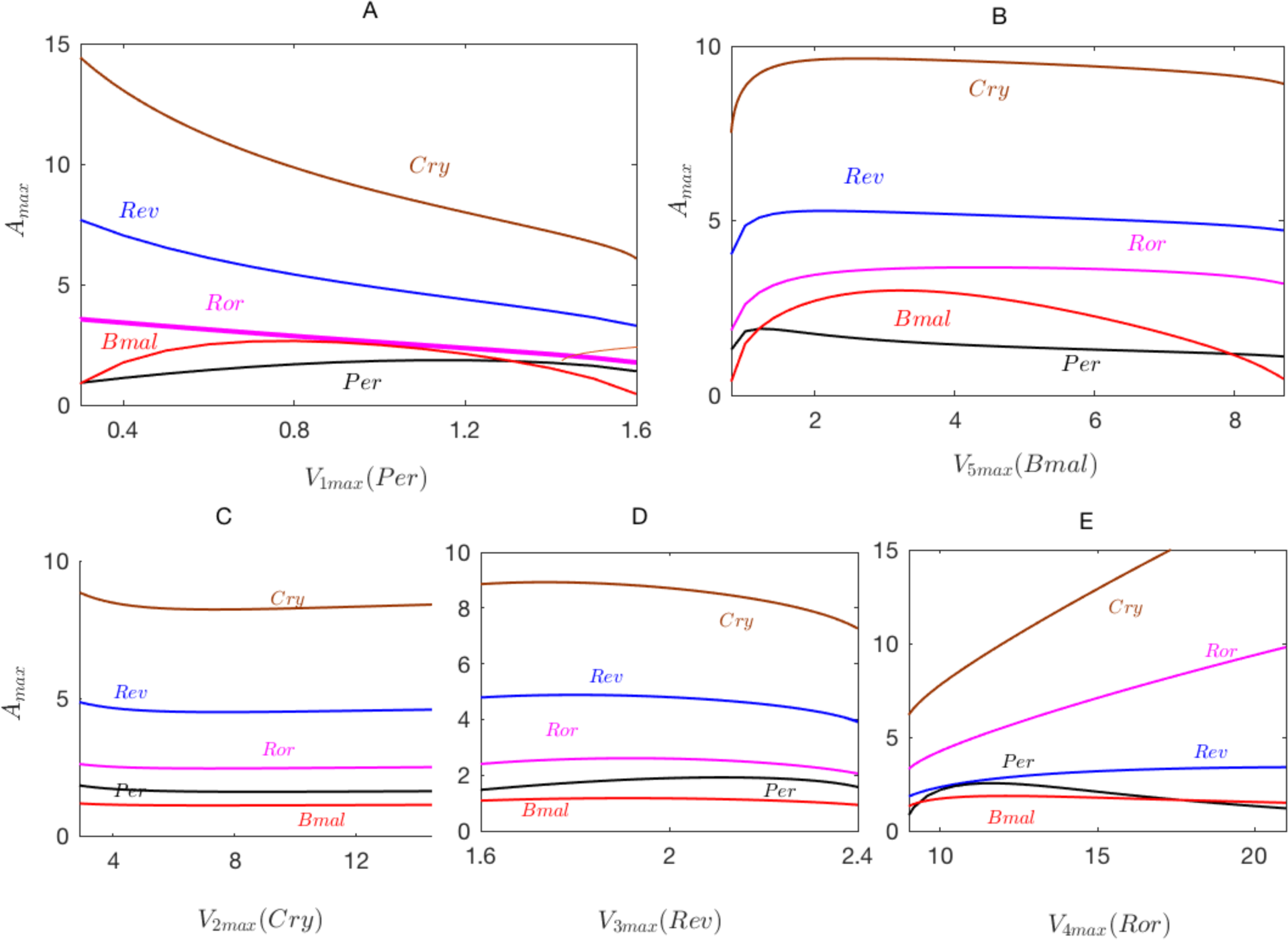
Dependence of the maximum amplitude of oscillations on the transcription rates V_*imax*_, *i* = 1..5. Other parameters ar fixed at WT values.

**Figure 7.**
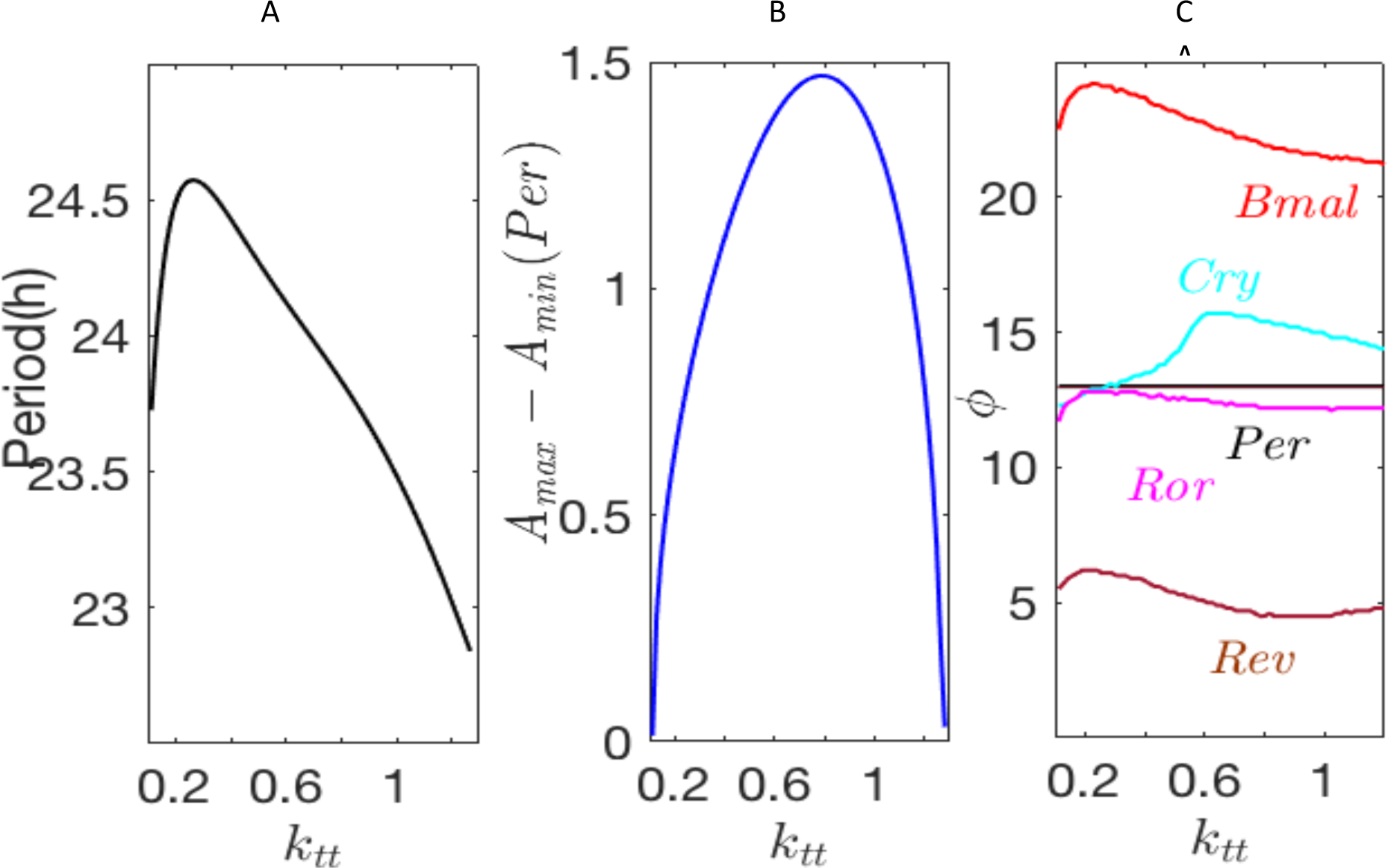
The effects of RAS signaling on the circadian clock network. Parameter *k*_*tt*_modulates the strength of transcriptional factor CLK:Bmal. A) Period. B) The difference between maximum and minimum amplitudes of *Per* oscillations. C) Phases of oscillation (relative to the phase of *Per*2 (black line) which is fixed at 13 h).

Keep in mind that the RBR loops are the union a REV loop (a negative-feedback transcription-translation loop) and a ROR loop (a positive-feedback transcription-translation loop). We shall make this distinction later.

### Numerical methods

The differential equations of Relogio’s model are provided in Suppl. Text S1. The variables of the model are listed in Suppl. Table S1, and WT (wild type) parameter values are provided in Suppl. Table S2. Numerical simulations of the model were carried out in Mathematica [32] and bifurcation diagrams were calculated using AUTO [33].

### Bifurcation diagrams

We use one- and two-parameter bifurcation diagrams to characterize the dependence of solutions of Relogio’s differential equations (S1-S19 in Suppl. Text S1) on the values of certain kinetic parameters in the model equations. In a one-parameter bifurcation diagram, the chosen kinetic parameter is plotted as the independent variable (the horizontal axis) and some indicator of the output of the model is plotted as the dependent variable (the vertical axis). Typically, the magnitude of a dynamic variable in the model is chosen as the dependent variable, in which cases the one-parameter bifurcation diagram shows the asymptotic behavior of this molecular component as a function of changes in the chosen kinetic parameter. In particular, the diagram shows how the steady-state value of the dynamic variable changes with the kinetic parameter and how the <u>amplitude</u> of any periodic solutions, bifurcating from the steady state, changes with the kinetic parameter. The diagram also indicates whether the steady states and periodic solutions are asymptotically stable or unstable.

We also use one-parameter bifurcation diagrams to indicate how the <u>period</u> of oscillatory solutions depends on a parameter value and how the <u>phases</u> of oscillatory components change, relative to one another, as the parameter value changes. In this way, we can use one-parameter bifurcation diagrams to succinctly summarize how the amplitude, period and phase of oscillations depend on the values of the kinetic parameters in a dynamical model of circadian rhythms.

We use two-parameter bifurcation diagrams to characterize the domain of oscillations on the plane of two chosen bifurcation parameters. We calculate period distributions inside the oscillatory domains and mark the regions of different periods with different symbols.

In complex mathematical models of biochemical reactions, involving multiple feedback loops, many different types of bifurcations and complex chaotic dynamics are expected [6]. In mammalian circadian systems, complex oscillations involving second and third harmonics have been observed [34]. Our simulations indicate that multiple harmonics and quasi-periodic dynamics are possible in Relogio’s model, Suppl. Fig. S1. In this work, we restrict ourselves to rhythmic dynamics generated by Hopf bifurcations.

## Results

### One-Parameter Bifurcation Diagrams

Previously, Relogio’a model was used successfully to simulate a variety of experimental conditions [16, 35, 36]. In this section, we extend simulations and predictions of Relogio’s model using bifurcation analysis, to show how principal characteristics of rhythmic dynamics (period, amplitude and phase) respond to continuous modulations of bifurcation parameters (primarily the maximum rates of expression of the core clock genes).

#### Modulations of PER expression

The one-parameter bifurcation diagram in Figure 2 shows how the amplitude of periodic solutions depend on the transcription rate of *Per* mRNA, *V*_1*max*_. All other parameters are fixed at WT values (Supplementary Table II). In the interval *V*_1*max*_ ∈ [0.142, 1.63], the model displays stable limit cycle oscillations with a circadian period. Notice that, at *V*_1*max*_ *≈* 0.25, there is a significant decrease of *Per* mRNA amplitude, due to multiple repressions exerted on the transcription of the *PER* gene. As reported in Ref. [16], there are small amplitude oscillations of long period (*T* ≈ 50 h) at small values of *V*_1*max*_. The two oscillatory domains in Figure 2 are separated by stable steady-state solutions. In Suppl. Fig. S2 we plot bifurcation diagrams of two other models of mammalian circadian rhythms and show that the small amplitude oscillations of long period are specific to Relogio’s model.

#### The period vs transcription rates

Rhythmic dynamics in a circadian network can be characterized by the period, amplitude, and phase of oscillations. The period is common to all variables, while the amplitudes and phases vary from one component to another. The solid lines in Figure 3 show that the oscillation period changes non-monotonically with changing transcription rates of the core clock genes, *V*_*imax*_,*i* = 1..5. An interesting feature in Figure 3 is that, as the transcription rates of the core clock genes increase, the period of the circadian rhythm eventually decreases, except for *ROR* gene transcription. The exceptional behavior of *ROR* indicates that the network reacts differently to *Ror* mRNA production than to increases of other mRNAs, presumably because ROR protein mediates the only positive feedback loop in the model.

#### Period vs mRNA degradation rates

Figure 4 shows how the period of oscillation changes with modulations of mRNA degradation rates. Period increases as the degradation rates of *Per* and *Cry* mRNAs increase above their WT values and decreases as the degradation rates of mRNAs in the RBR loops increase above their WT values. In the case of *Rev*, the period change is limited to a narrow interval [23h, 23.7h], whereas the period change is much more pronounced for variations in the degradation rates of *Ror* and *Bmal*. An interesting feature for *Bmal* is that the period is no longer “circadian” for small values of *d*_*y5*_.

#### Phase relations vs transcription rates

We define the phase of oscillations as the time when mRNA level of a core clock gene reaches its maximum level, relative to the phase of a reference mRNA. Figure 5 shows that phases do not change significantly with modulations of transcription rates. In general, the phase of *Cry* is most sensitive to changes of the transcription rates. The phase of *Bmal* changes moderately with changes of transcription rates of the RBR loop. With changing rate of transcription of the *PER* gene, the phases of *Ror* and *Bmal* are locked, but the phases of *Rev* and *Cry* change slightly. With changing rate of transcription of the *REV* gene, the phase of *Ror* is locked, but the phases of *Bmal, Rev* and *Cry* change slightly.

#### Amplitudes vs transcription rates

Figure 6 shows the changes of the maximum amplitudes of oscillations, *A*_*max*_,with the change of transcription rates. The amplitudes of *Per* and *Ror* do not change significantly in Figure 6, indicating robustness of these oscillatory variables. By contrast, the amplitude of *Ror* is sensitive to modulations of transcription rates. The amplitude of *Cry* is sensitive to *V*_2*max*_(*cry*), as well as to modulations of the positive feedback loop *V*_*4max*_(Ror) and *V*_*5max*_(*Bmal*). The amplitude of *Bmal* decreases at large values of transcription rates.

#### Modulations of all transcription rates

Experimental data on *α*-amanitin-treated and actinomycin D-treated fibroblast cells reported by Dibner et al. [37] showed that mammalian circadian oscillators are resilient to large fluctuations of transcription rates. Indeed, a general reduction of transcription rates of all mRNAs leads to oscillations with reduced period and amplitude. Interestingly, circadian rhythm models based on a delayed negative feedbacks loop (e.g., the Goodwin [8] and Leloup-Goldbeter [13] models) display similar features when the transcription rates of all mRNAs are reduced. We found that Relogio’s model displays similar modulations of period and amplitude, when all transcription rates are down-regulated. However, period and amplitude are initially increased and then reduced if the transcription rates are increased from WT values (see Suppl. Fig. S3).

#### Interactions between the circadian clock and RAS/MAPK signaling networks

The dynamics of period, amplitude, and phases, obtained by bifurcation analysis can be used for practical purposes. Experimental data about the interactions between RAS/MAPK and the core clock network can be simulated using Relogio model. The RAS/MAPK signaling pathway may disrupt normal functioning of the core clock network by modulating the circadian period [35, 36]. It is known that changes in *RAS* expression alter directly the dynamics of *BMAL* expression [35]. Therefore, perturbation of the core clock network by RAS signaling has been studied by introducing a parameter *k*_*tt*_, which modulates the strength of the transcription factor CLK:BMAL (variable *x*_1_) for the expression of every clock gene [35, 36]. We refer to the S1 text of Ref. [36] for the definition of *k*_*tt*_. In Figure 7 we show how period, *Per*’s amplitude, and phases of oscillation change with *k*_*tt*_. The WT value of *k*_*tt*_ *=* 1.

Notice qualitative similarities between the periods in Figure 3A and Figure 7A, as well as between *Per*’s amplitude in Figure 6A and Figure 7B. Recall that Figure 3A and Figure 6A show the network’s response to the modulation of *Per’*s transcription rate, whereas Figure 7A-B show the RAS/MAPK pathway’s interference on *Bmal’*s transcriptional activation. The period change in Figure 7A is similar to the period change in Figure 3A where *V*_1*max*_ (transcription rate of *Per*) is modulated, suggesting that BMAL activates strongly the transcription rate of *Per* mRNA. Interestingly, the changes of oscillation phases with *k*_*tt*_ are similar to phase changes in Figure 5E, where *V*_5*max*_ (the transcription rate of *Bmal*) is modulated.

Figure 7 suggests that by monitoring the changes in the expression patterns of a given clock gene caused by modulations from other pathways in the cell, qualitative changes in the period, amplitudes, and phases of oscillations can be inferred from bifurcation diagrams in Figure 3-6. As an example, an antisense transcript, *Per*2*AS*, of the *PER*2 gene may depress the transcription rate of its sense counterpart, *Per*2 [35]. Figure 6A shows how the level of *Per*2 will be reduced when *V*_1*max*_ is depressed. Furthermore, according to Figure 3A, the period of oscillation is expected to increase, but according to Figure 5A, the phases of oscillations are expected to change slowly. Detailed simulations of a mathematical model describing molecular interactions between sense and antisense transcripts are in agreement with these assessments [35].

It was reported that the effects of RAS modulations on the core clock network can be described qualitatively by modulations of a single parameter, *k*_*t*2_ [35]. Supplementary Fig. S4 shows the changes of period, amplitude (*Per*), and phases of oscillations with changing values of *k*_*t*2_ (Eq. (S12)). The main difference between modulations by *k*_*tt*_ and *k*_*t*2_ is in changes of *Cry* phase.

### Two-Parameter Bifurcation Diagrams

Although many different mathematical models have been proposed for mammalian circadian rhythms [14], to the best of our knowledge, detailed parameter “portraits” of these models are scarce, despite the importance of the stability and robustness of circadian rhythms against simultaneous perturbations of multiple genes and feedback loops. In this section, we study the stability of oscillatory domains in Relogio’s model with simultaneous modulations of a pair of parameters specifying interaction strengths between selected genes and feedback loops.

#### Interactions between Bmal and Per

Figure 8 shows a two-parameter bifurcation diagram of Relogio’s model for *V*_1*max*_ (*Per*) and *V*_5*max*_ (*Bmal*), transcription rates of a repressor and activator, respectively. The solid lines in Figure 8 are the loci of the four Hopf bifurcation points in Figure 2. Stable oscillations are found inside the solid curves. We marked the large oscillatory domain in Figure 8 with symbols of different colors to show how the period of oscillation depends on the values of these two parameters. In the simulations, we fixed all other parameters at WT values. Inside the small ellipse shown in Figure 8, we found the small-amplitude, long-period oscillations shown in Figure 2 by red symbols. The horizontal dashed line in Figure 8 shows that as *V*_1*max*_ increases, oscillations are replaced by stable, steady state solutions at *V*_1*max*_ ≈ 1.63. The oscillations outside the black lines are slowly damped. Although we do not focus on damped oscillations in this work, they may well be relevant to the generation of robust circadian rhythms by global synchronization of populations of coupled damped oscillators [38].

**Figure 8.**
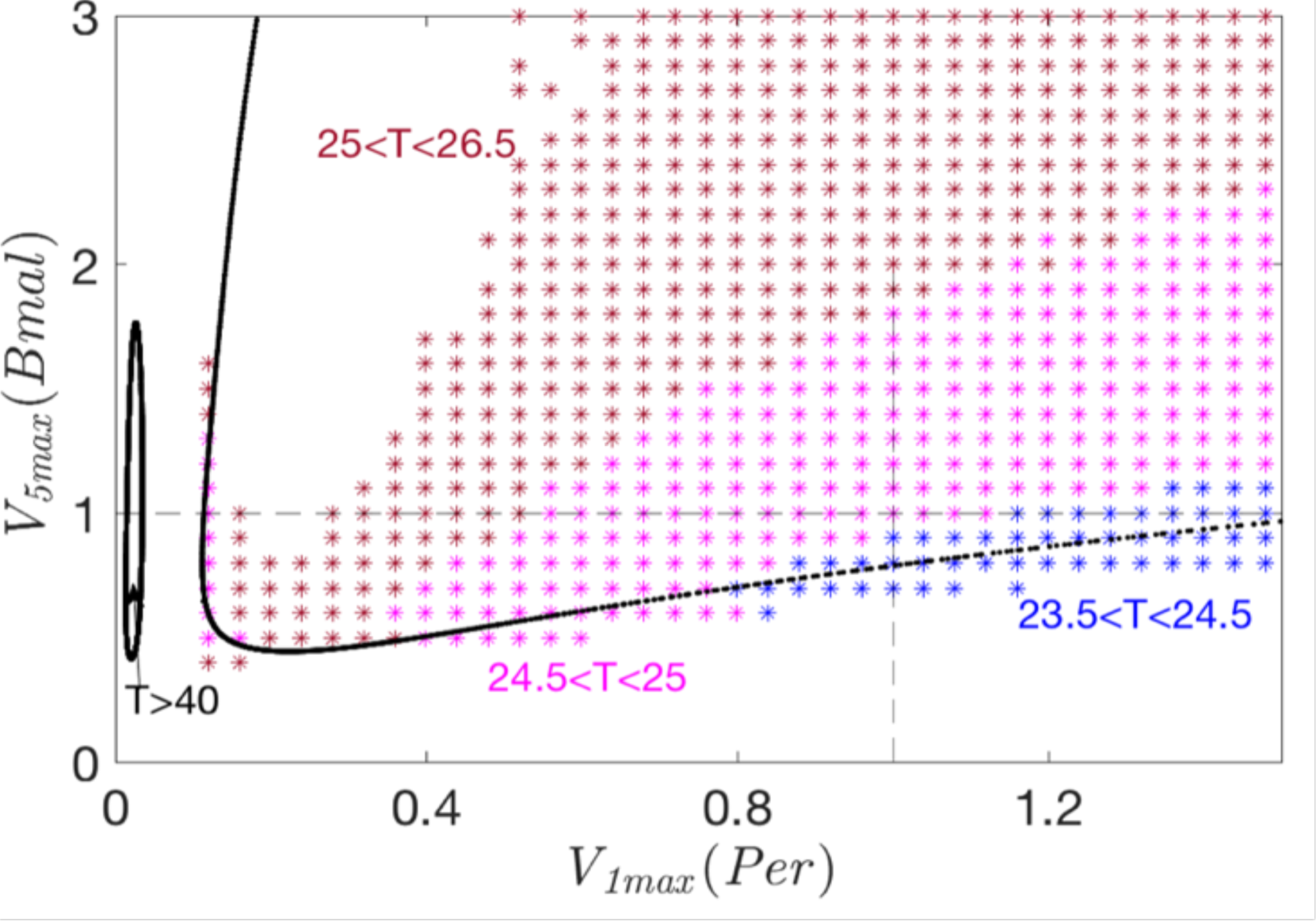
A two-parameter bifurcation diagram on the (*V*_1*max*_, *V*_5*max*_) parameter plane. The symbols indicate the period of oscillations at a given location on the parameter plane. Dashed lines mark the WT values of *V*_1*max*_ and *V*_5*max*_ Slow oscillations inside the small ellipse near *V*_1*max*_≈ 0.05 correspond to the oscillations shown in Figure 2 by red symbols. In the white area inside the large domain, the period is greater than 26.5 h.

#### Period locking in the PC loop

Figure 9A shows a two-parameter bifurcation diagram on the parameter plane (*V*_1*max*_, *V*_2*max*_), the transcription rates of *Per* and *Cry.* The solid red curves in Figure 9A show the loci of Hopf bifurcation points *HB*_1_ and *HB*_4_ in Figure 2. Symbols of different colors denote the period of oscillations. An interesting feature of Relogio’s model is that the period of oscillation can be locked within a narrow interval, if either *V*_1*max*_ or *V*_2*max*_ is modulated alone. This can be seen, for example, by following the vertical dashed line in Figure 9, above the horizontal dashed line. In the case of *V*_1*max*_’s modulation, *V*_2*max*_ must be set to a lower value than its WT value, in order to observe period-locking over a wide range of *V*_1*max*_ values.

**Figure 9.**
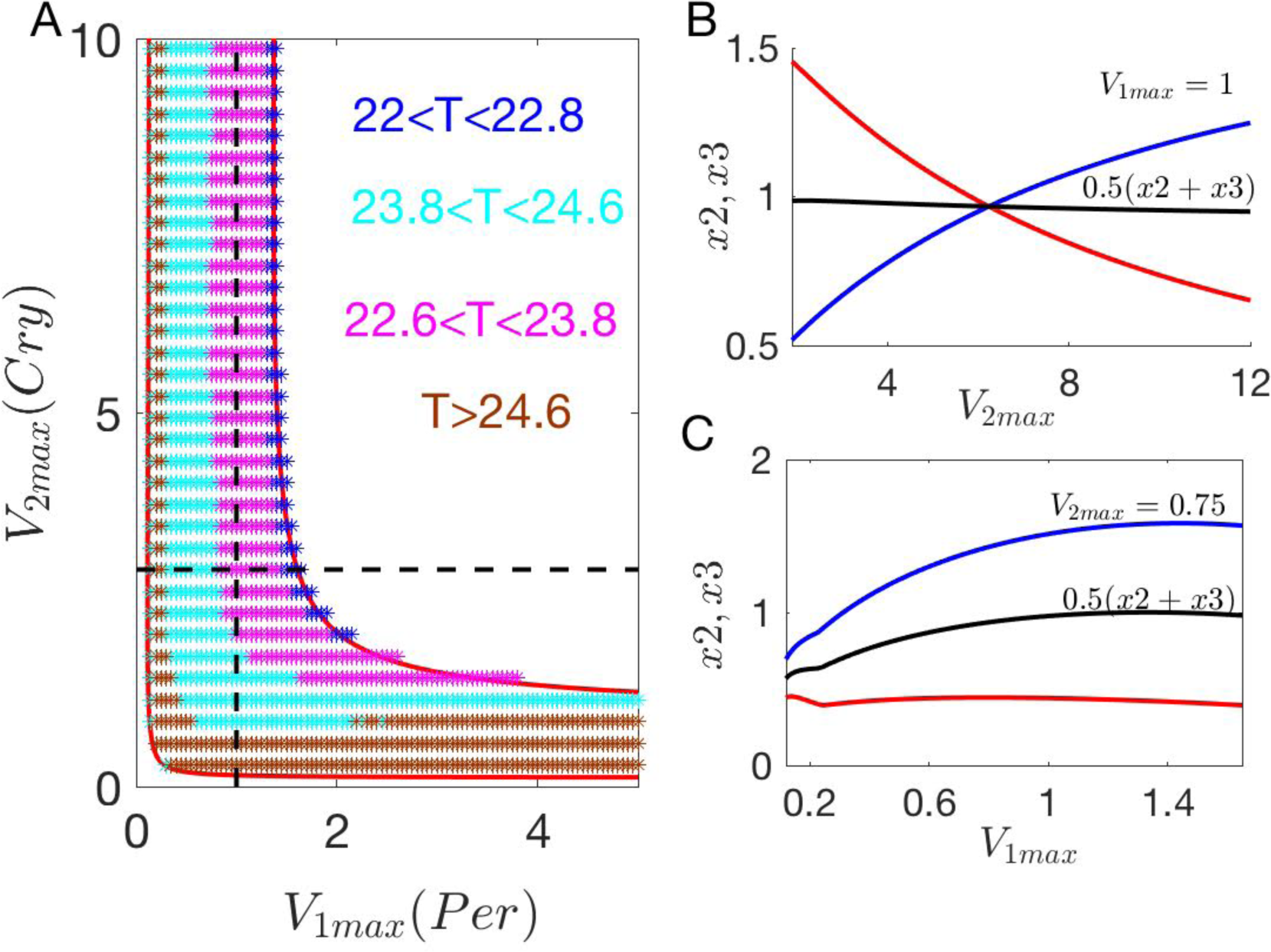
The dynamics of the PC loop. (A) Two-parameter bifurcation diagram on (*V*_1*max*_, *V*_2*max*_), all other parameters are fixed at their WT values. Symbols mark the period of oscillation. When *V*_1*max*_*=* 1 (WT value), the period of oscillations is insensitive to changes of 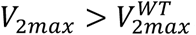 Also, when *V*_2*max*_≤ 0.75, the period is insensitive to the changes of *V*_1*max*_ Dashed lines mark WT values of *V*_1*max*_and *V*_2*max*_ (B, C) The maximum amplitudes of *x*2(blue), *x*3(red), and *PC* (black) vs *V*_2*max*_, when *V*_1*max*_is fixed at 1, and vs *V*_1*max*_, when *V*_2*max*_is fixed at 0.75.

Figures 9B-C provide insight into period-locking in Relogio’s model. These panels show how the maximum amplitudes of *x*2 (nuclear phosphorylated PER/CRY complex), *x*3 (nuclear unphosphorylated PER/CRY complex) and *PC = x*2 + *x*3 (*PC* pool) change with changes of *V*_1*max*_ and *V*_2*max*_. With respect to increasing values of *V*_2*max*_ (panel B), *x*2 level increases (blue lines) and *x*3 level decreases (red lines), so that *PC* level is compensated (black lines). Consequently, the repression of core-clock genes by *PC* changes little with increasing *V*_2*max*_. Thus, slow changes of *PC* restrain the period within a narrow interval. Figure 9C shows a similar compensation of *PC*, when *V*_1*max*_ is modulated, for *V*_2*max*_ *=* 0.75.

#### Interactions between PC and REV loops

In Figure 10 we explore the interplay of the negative-feedback transcription-translation loops, PC and REV. To this end, we introduce a new parameter *α*_*pc*_, *PC = α*_*PC*_(*x*2 + *x*3), to modulate the activity of PER/CRY in the nucleus: *α*_*PC*_ is used to boost PC activity up or turn down relative to the WT value, *α*_*PC*_ = 1. The blue curve in Figure 10A shows the locus of *HB*_1_ points (see Figure 2) on the (V_Smax,_, *α*_*PC*_) parameter plane. The periods of oscillations for 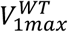 in different regions are marked by different symbols. Note that some of the symbols are outside of the blue lines because of 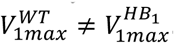 All other parameters are fixed at WT values. Interestingly, the black symbols obtained by simulations reveal that there is another domain of oscillations on (*V*_3*max*,_ *α*_*PC*_), where the period is *T* > 32 h. Because nuclear protein REV_N_ directly represses *Cry*, the PC loop is down regulated with the increase of *V*_3*max*_. As a result, REV_N_ releases *PC*’s repression of *Bmal,* allowing the RBR loops to generate slow oscillations shown in Figure 10A.

**Figure 10.**
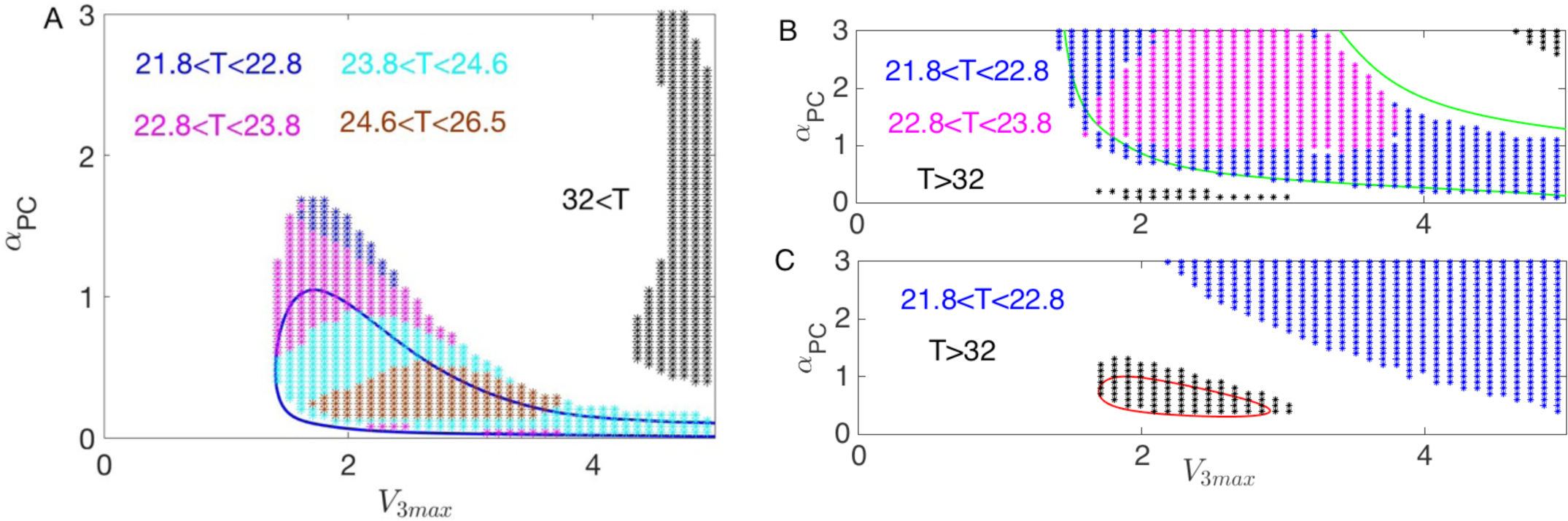
Interactions between the PC and REV loops, as revealed by two-parameter bifurcation diagrams. (A) The blue lines show the loci of *HB*_1_in Figure 2. The symbols of different colors denote the oscillatory periods in different regions. Black symbols at V_3*max*_> 4 show a domain of slow oscillations. (B) Green lines show the continuation of *HB*_2_in Figure 2. (C) Red lines show the continuation of *HB*_3_ in Figure 2.

In Figure 10B we show the continuation of the bifurcation point *HB*_2_ in Figure 2. Blue and purple symbols mark oscillation periods at 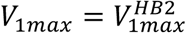 Because 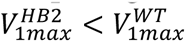, the amplitude of *Per* oscillations is low. Black symbols in Figure 10B show that besides the oscillatory domain inside the loci of *HB*_2_, there are two separate domains of slow oscillations on (*V*_3*max*_, *α*_*PC*_). As the lower domain of slow oscillations persist at *α*_*PC*_ ≈ 0, here the strength of PC loop is negligible compared to REV’s repression. As for the upper slow domain, though *α*_*PC*_ is strong in this region, *V*_3*max*_ is even stronger; hence, the RBR loops can independently generate slow oscillations.

The red curve in Figure 10C show the continuation of the bifurcation point *HB*_3_ in Figure 2. The amplitude of *Per* oscillations is small because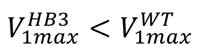. Interestingly, at larger values of *V*_3*max*_ and/or *α*_*PC*_, oscillations with a circadian period emerge in Figure 10C, blue symbols. However, the amplitude of *Per* oscillations is again small due to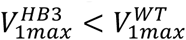.

We note that the loci of the bifurcation point *HB*_4_ has similar features as the loci of the bifurcation point *HB*_3_. In Suppl. Figs. S5-S6, we explore interactions between the PC and ROR loops, and the REV and ROR loops, respectively. Taken together, our two-parameter bifurcation diagrams show that interlocked feedback loops generate multiple domains of oscillations in Relogio’s model.

### The Role of the REV Feedback Loop

Recent ChIP data show that the REV loop plays a critical role in the core clock [39]. In addition, WT levels of RevErb*α* and RevErb*β* protect circadian clock and normal metabolic functions, while the double knockouts lead to clear non-rhythmic phenotypes [40, 41]. Previous theoretical work by Relogio et al. [16] showed that, in the intertwined dynamics of their model, the REV loop plays a dominant role in balancing positive and negative interactions.

The goal of this subsection is to uncover the critical role of the REV negative-feedback loop in the generation of sustained oscillations, by clamping certain genes near their mean values [14, 16], which allows us to fix some state variables at constant levels, thereby reducing the number of ODEs in the model. For example, when the state variable *x*_6_ (nuclear protein, ROR_N_) is fixed near its mean WT level (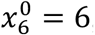, which means that *Ror* mRNA was clamped near its mean expression level, *Ror* = 3.8), the network is rhythmic. Furthermore, if we clamp *PC* (*x*_4_ + *x*_3_) near its mean level (*PC* ≈ 1.7, which means that *Per* and *Cry* mRNAs were clamped near their mean expression levels, *Per* = 1.1 and *Cry* = 4.8), the system is still rhythmic [16], see Suppl. Fig. S7. However, if *PC* is clamped at more than +20% from its mean level, the system becomes non-oscillatory. Note that with the double clamping of *PC* and *x*_6_, the original model, composed of 19 ODE’s, is reduced to 7 ODE’s describing the state variables in the REV-BMAL loop only (Eqs. (S21-27)). Although such an overly simplified model cannot serve for modeling experimental data, it is helpful for uncovering the role of the REV loop.

For appropriately chosen fixed values of *PC* and *x*_6_, Eqs. (S21-27) display oscillatory dynamics. In Figure 11, we plot a bifurcation diagram on (*V*_3*max*_, *V*_5*max*_), for 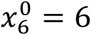 and *PC*^0^ *=* 2.04. The black line shows the locus of Hopf bifurcation points in the reduced model. For the sake of comparison, we plot in Figure 11 (dashed green line) the locus of Hopf bifurcation points *HB*_1_ in the full model. The oscillatory domains in the reduced and full models exhibit similar shapes. Moreover, as Suppl. Fig. S8 shows, the REV-BMAL loop can drive circadian oscillations independently of the PC and ROR loops.

**Figure 11.**
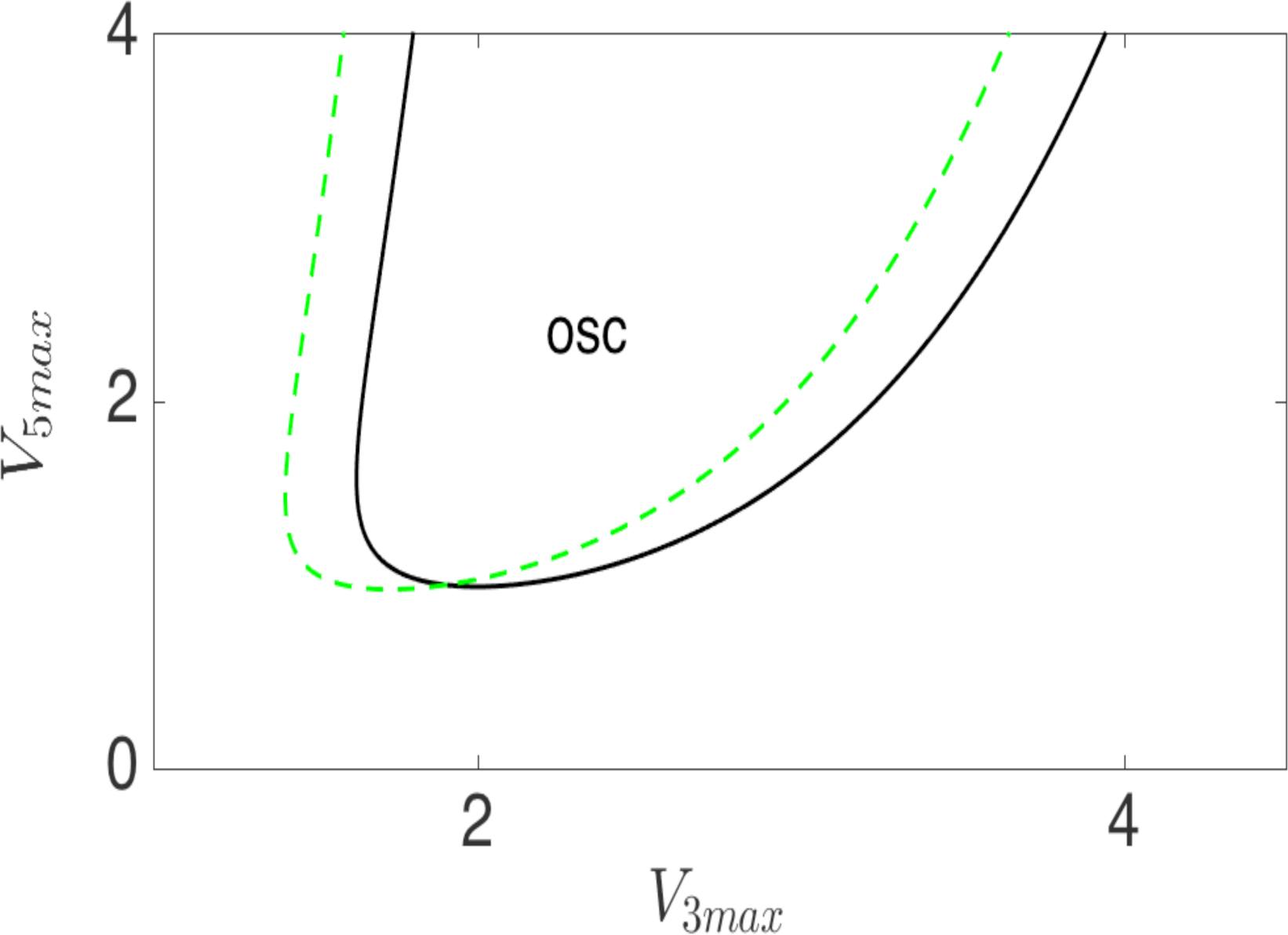
Two-parameter bifurcation diagram on the (V_3*max*_, *V*_5*max*_) plane. The solid black curve shows the locus o Hopf bifurcation points in the reduced seven-variable model (Eqs. (S21-27)). The dashed green curve shows the locu of *HB*_1_(Figure 2) in the full Relogio model. Other parameters of the reduced model are the same as Relogio’s W parameters, and 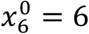 and *PC*^^^ *=* 2.04.

*Constitutive expression of* REV. Previous experimental studies of the circadian clock in mammalian cells have investigated by effects of constitutively expressing the core clock genes *CRY*, *BMAL*, and *PER*1 [42-44]. It was reported in Ref. [16] that when REV is constitutively expressed (when REV_N_ level is held constant at 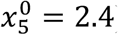), the circadian rhythm is lost. (Note that when 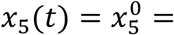, Relogio’s model reduces to a system of 16 ODEs.) If 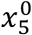 is further increased, the reduced model recovers oscillatory dynamics [16], because of the effects of 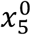 on the balance of positive and negative feedbacks in the system. The goal of this subsection is to use two-parameter bifurcation diagrams to reveal dynamic transformations in the network, when REV is constitutively expressed.

First of all, to illustrate the effects of constitutively expressed REV, we choose to modulate two parameters that control the strength of the PC loop (Figure 12), namely, *V*_*5max*_, the maximum rate of expression of *Bmal* mRNA, and *c*, the Hill exponent for *PC*’s repression of *Per* mRNA synthesis in Eq. (S11). The dashed green curve in Figure 12 shows the locus of *HB*_1_ and *HB*_2_ (see Figure 2) on the(*c*, *V*_5*max*_)parameter plane. Inside the dashed green line, Relogio’s model displays oscillatory dynamics with a circadian period. Dashed black lines in Figure 12 mark WT values of *c* and *V*_5*max*_ When 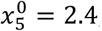, the reduced model displays a steady-state solution. We found that if the transcription rate of *Bmal* is reduced from its WT value, an oscillatory domain emerges, shown by the blue solid line in Figure 12. In this domain the period (*T* > 35 h) is larger than circadian. Because the domain of slow oscillations is persistent at *c* = 0, the role of the PC loop can become negligible in this region.

**Figure 12.**
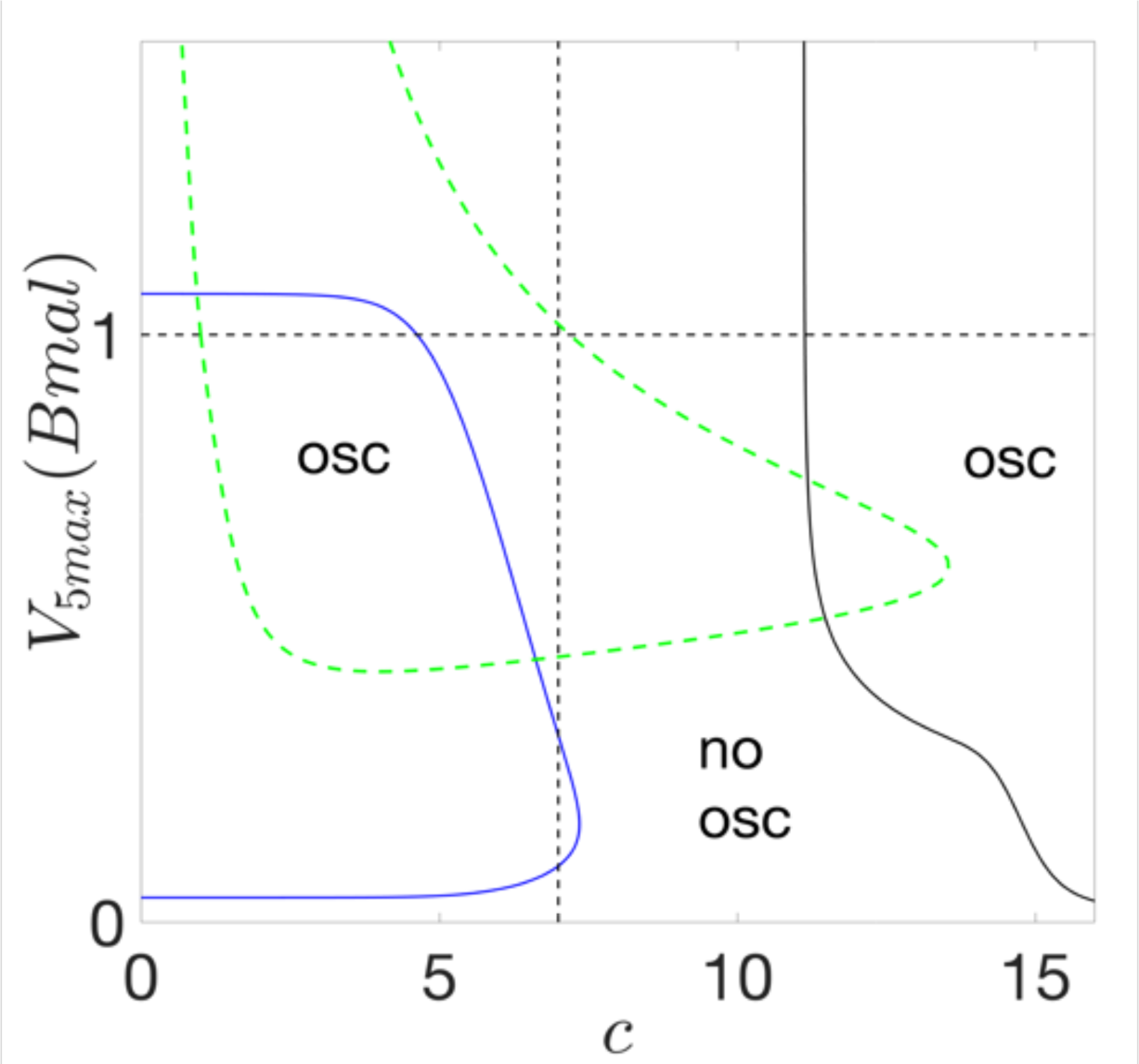
Two-parameter bifurcation diagram. The dashed green curve shows the locus of Hopf bifurcation points *HB*_1_ and *HB*_2_in Figure 2. If the level of REV is fixed at 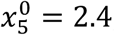, the system becomes non-oscillatory. However, at different values of the parameter *c* in Eq. S11, two independent oscillatory domains emerge. The blue solid curve bounds a domain of slow oscillations (period > 35 h) at small *c*. The black curve shows a boundary of oscillations at larger *c*, where the period of oscillations is ∼20 h.

There is yet another oscillatory domain, shown by the solid black line in Figure 12, at larger values of *c*. Because at larger values of *c* the repression of *PER* gene expression by PC is stronger, the PC loop recovers its intensity in this domain. Here the period of the oscillations (∼20 h) is smaller than circadian. These oscillations persist even if the BMAL-ROR loop becomes constitutive 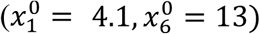

The recovery of oscillations dampened by constitutive expression of REV suggests that the balance of positive and negative regulations in the network of interlocked loops is critical for maintaining the circadian rhythm. When REV is constitutively overexpressed 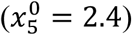, the synergy of interlocked loops is lost and the balance of positive and negative regulations is shifted towards the dominance of positive regulation. To recover oscillatory dynamics, either the intensity of the positive feedback needs to be weakened (decreasing *V*_5*max*_) or the intensity of the negative feedback needs to be strengthened (increasing *c*). Thus, the critical role of the REV loop in Relogio’s model is in interfacing and synergizing positive and negative regulations of the multiple interlocked loops.

*Deformation of oscillatory domains.* The core-clock network interacts with other pathways in the cell and responds to global regulatory signals in the organism. Depending on how an external signal modulates the core clock network, rhythmic dynamics can be distorted; hence, the underlying oscillatory domains can be deformed. Such dynamic transformations of the core clock network can be studied by bifurcation analysis.

In this subsection, we illustrate deformations of an oscillatory domain in Relogio’s model when the REV loop is modulated by internetwork interactions, for example, by interactions with the cell cycle network. We consider two hypothetical modulations, both acting on the variable *x*_1_ (nuclear protein REV_N_).

<u>Hypothetical modulation #1</u>: we assume in Eq. (1) that as a result of external perturbations, the production of nuclear protein REV_N_ can be increased or decreased by a factor *β*_*x*5_,

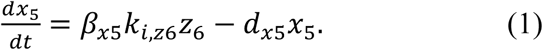

<u>Hypothetical modulation #2</u>: we assume in Eq. (2) simultaneous perturbation of the rates of production and degradation of REV_N_, so that the parameter *β*_*x*5_ now modulates the time scale of *x*_5_,

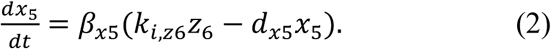

In both cases, the WT value of *β*_*x*5_ = 1.

In the first case, we replaced the ODE for *x*_5_ (Eq. (S6)) in Relogio’s model with Eq. (1). Figure 13 shows the two-parameter bifurcation diagram on (*V*_1*max*_, *β*_*x*5_). The green dashed curve shows the locus of Hopf bifurcation points *HB*_1_ and *HB*_2_ in Figure 2; the mauve dashed line shows the locus of Hopf bifurcation points *HB*_3_ and *HB*_4_ in Figure 2. Inside the mauve and green lines, there are two separate oscillatory domains with slow and circadian oscillations, respectively. Both domains shrink and disappear with changing values of *β*_*x*5_.

**Figure 13.**
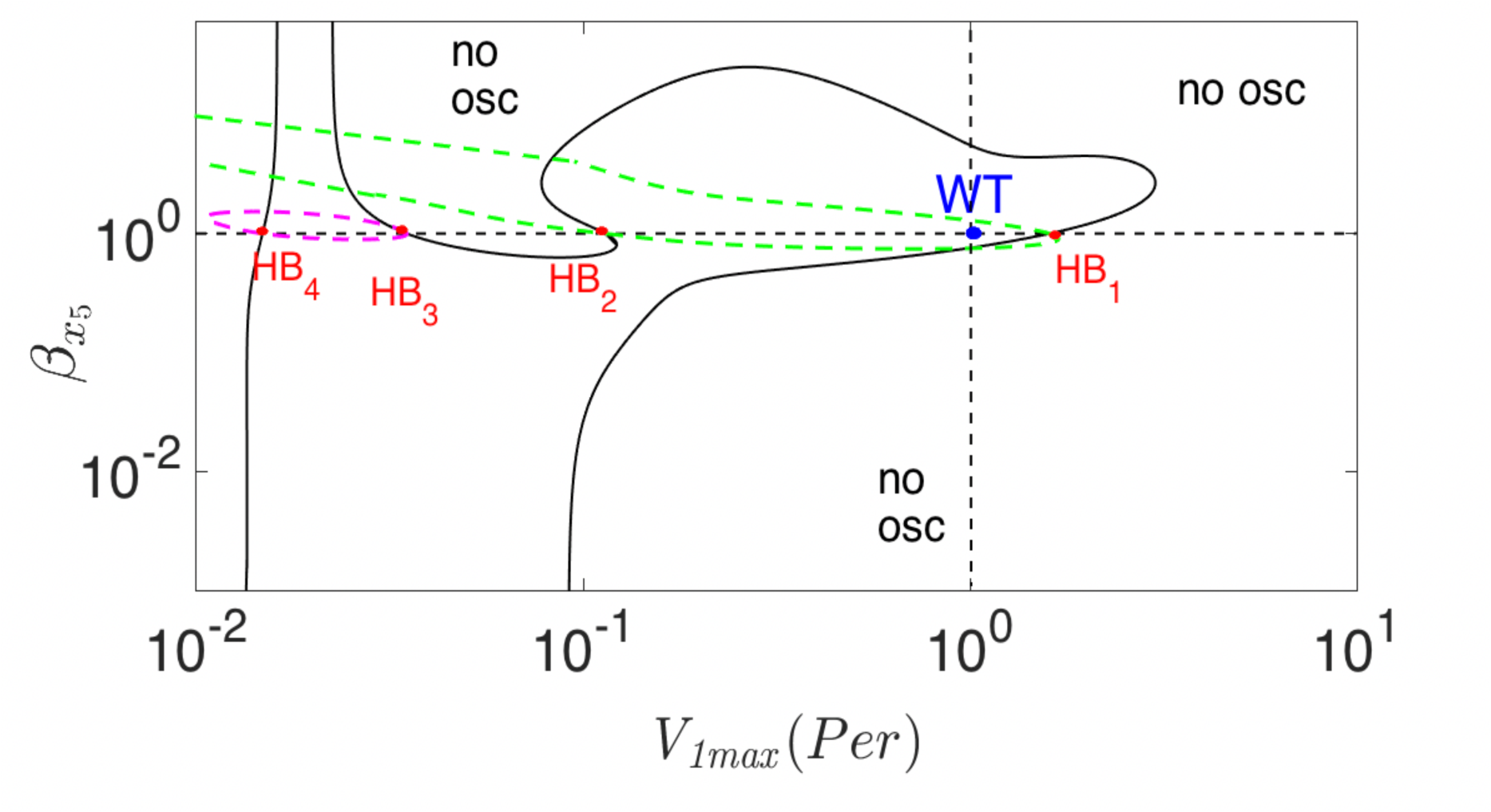
Modulations of the REV feedback loop. *β*_*x*5_is a parameter introduced in Eq. (1) and Eq. (2) to modulate, respectively, either the level of *x*_5_(nuclear protein REV_N_) or the time-scale of *x*_5_. Dashed green and mauve lines are obtained using Eq. (1). *HB*_*i*_(*i* = 1..4) mark the Hopf bifurcation points in Figure 2. Solid black lines are obtained using Eq. (2). Inside the solid lines, the system displays oscillatory dynamics.

Next, we replace Eq. (S6) with Eq. (2). The two black, solid curves in Figure 13 show the loci of the Hopf bifurcation points *HB*_*i*_, *i* = 1..4, in Figure 2. In contrast to the mauve and green lines, there is a single domain inside the black curves. At small values of *β*_*x*5_ and *V*_1*max*_, or at large *β*_*x*5_ but small *V*_1*max*_, the black lines are almost parallel to each other. In these regions, there are only two intersections of the oscillatory domain with a horizontal line. Clearly, the domain of oscillation is larger for the solid curves; remarkably, its size increases with an increase of *β*_*x*5_, above the intersections of dashed lines. Note that emergent domains of oscillations are also possible in Figure 13, similar to the slow oscillations shown in Figure 10 by black symbols.

In order to effectively synchronize cellular processes, the core clock network must be resilient to various perturbations. As Figure 13 shows, robustness of the rhythm in the network against parametric modulations can be characterized by deformation of an oscillatory domain. The dashed lines in Figure 13 show that oscillatory domains may shrink if the perturbation modulates the production of REV_N_ in Eq. (1). But the solid lines in Figure 13 show that if the processes counterbalancing such a perturbation can be activated, as in Eq. (2), the size of a circadian domain can be increased. As enlargement of an oscillatory domain typically implies enhancement of the rhythm’s robustness, modulations from other pathways in the cell may strengthen the synergy of the feedback loops. Interestingly, the interactions between core clock and cell cycle networks is interfaced by the REV loop [23, 25] which plays a critical role in the synergy of the interlocked loop.

## Discussion

Comprehensive mathematical models of fundamental biological processes, such as circadian rhythms or cell cycle controls, play important roles in molecular systems biology. Bifurcation analysis is a powerful tool for qualitative characterization of these mathematical models [21, 45]. For a model with many variables and parameters, bifurcation analysis can be very informative and also very challenging. In this paper, we have used bifurcation analysis to decipher the roles of interlocked feedback loops in a model of the mammalian circadian-clock network, namely, a model proposed by Relogio et al. [16].

Our bifurcation diagrams show how the period, amplitudes and phases of the mammalian circadian rhythm depend on levels of expression of the core clock genes and other principal parameters in Relogio’s model. In principle, any of these predictions can be tested experimentally by varying the rates of expression of core clock genes up or down from WT values by standard techniques of molecular genetics. If we can control the expression of two or more of these genes simultaneously, then we can test the model’s predictions—contained in two-parameter bifurcation diagrams (e.g., Figures 8 and 9)—of the effects of interactions between pairs of genes. Relogio et al. [16] have already considered some of these predictions in light of experimental results, but there is much that can and should be done in the future by way of experimental testing of the interactions of the feedback loops of the mammalian circadian clock.

Admittedly, our current understanding of the mammalian circadian-clock network is incomplete. Bioinformatics studies have revealed more than 40 genes that directly interact with the core clock genes [46]. Together, these known and yet-to-be-discovered clock genes will form an extended clock network. As the extended network comes into focus and detailed mathematical models are developed, the consequences of the feedback and feed-forward loops in the network will become ever harder to recognize and understand. We propose that one- and two-parameter bifurcation diagrams, as presented in this work, will be essential in revealing the dynamic changes of period, amplitude and phase of oscillations, as well as deformations of oscillatory domains in the circadian clock network, in dependence on rates of gene expression and other crucial quantitative features of the genetic control system (e.g., protein stability, feedback intensities, etc.). The parameters whose modulations cause qualitative changes in network dynamics will identify genetic components and feedback loops that may be susceptible to pharmaceutical interventions, in cases of circadian dysrhythmias.

With the present array of circadian-clock models, we can already see how such analytical tools may be useful. Depending on how an oscillatory model is built and parameterized, the mechanisms generating oscillations can be different. Unlike previous models of circadian oscillations, which rely chiefly on a single negative-feedback transcription-translation loop (the classical Goodwin-type mechanism of oscillation [12]), the circadian rhythm in Relogio’s model is driven by three synergetic feedback loops [16]. Our bifurcation analysis shows that (similar to the Goodwin mechanism) the PC loop can display a single Hopf bifurcation point as the vertical line in Figure 9 shows. The persistence of oscillations with the increase of *Cry*’s transcription rate is consistent with the *Cry* overexpression phenotype retaining the circadian rhythmicity. On the other hand, a bifurcation diagram of the REV-BMAL-ROR (RBR) loops typically displays a second Hopf bifurcation point as the horizontal line in Figure 8 shows, due to the positive feedback exerted on *REV*. Such behavior is consistent with the reported aperiodic dynamics for gene overexpression in the RBR loops [16]. Thus, the mechanism of interlocking loops explains overexpression phenotypes in mice, by combining contrasting dynamics: *Cry* overexpression retains rhythmicity, gene overexpression in the RBR loops loses rhythmicity [16].

In Relogio’s model, the RBR loops play a special role. Oscillations persist if the PC loop is non-oscillatory (*x*_4_ *= const*, *x*_3_ *= const*), but they damp out if the RBR loops are non-oscillatory. It was reported that when *REV* is constitutively expressed 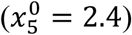, WT oscillations are damped out [16]. However, we find that, even with constitutively overexpressed *REV* 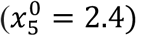, oscillations are possible at different values of the parameter *c* (see Figure 12). These oscillations are generated by the PC and RBR loops independently in two separate domains. In the first domain, oscillations are slow and persistent even if the PC loop is constitutive. In the second domain, oscillations are rapid and persistent even if the BMAL and ROR loops are constitutive. Thus, it is the REV loop that enables the synergy of positive and negative feedback loops to generate robust circadian rhythms in Relogio’s model.

Our bifurcation diagrams suggest that the mammalian core clock network operates by balancing the activities of interlocked feedback loops and if the balance is lost there are multiple mechanisms for generating rhythmic dynamics. In Relogio’s model, if circadian rhythms are lost due to modulations of the PC loop, then the RBR loops can generate stable, slow oscillations. Interestingly, in a different model of circadian rhythms [12], when the transcription rates of *Per*1 and *Per*2 are zero, the system becomes non-rhythmic if all other parameters are at their WT values. However, if REV is overexpressed, stable, fast oscillations can be generated (Suppl. Fig. S9). Therefore, a critical question arises from predictions of the Relogio and Mirsky et al models as to whether different mechanisms of oscillations can be activated in experimental systems. We suppose that the question can be answered by an experiment where transcription rates of multiple repressor genes can be modulated. The knock down of repressors in the PC loop, e.g., *Per*1 and *Per*2, can induce arrhythmic dynamics [1]. Then if REV is overexpressed, for example by activating CLOCK/BMAL expression or by constitutive REV overexpression [16], the circadian rhythm can be revived due to the existence of multiple oscillatory mechanisms in the network, as mathematical models of interlocked feedback loops predict.

In extended models, the circadian core clock network is integrated to other pathways in the cell, which can themselves display autonomous oscillation [23, 25, 27, 32, 36]. A plethora of oscillatory mechanisms and domains of oscillations are possible in such models. Intuitively, the size, shape, and number of oscillatory domains can be indicators of robustness of rhythmic dynamics. But how the interplay of different oscillatory mechanisms is regulated and how complex networks integrate oscillatory domains are still unknown and remain major goals of future studies [13].

## Acknowledgements

Financial Assistance from the Department of Biological Sciences and the College of Science at Virginia Tech is gratefully acknowledged. DB was partially supported by a grant from the Mongolian Foundation for Science and Technology. DB is thankful to Dr. Angela Relogio for explaining the results published in [16]. DB is also grateful to Prof. Kh. Namsrai for stimulating discussions.

## Author Contributions

Designed research (DB) and (JJT); performed research (DB); visualization (DB); wrote the manuscript (DB) and (JJT).

## References

1. Reppert SM, Weaver DR. Coordination of circadian timing in mammals. Nature. 2002;418(6901):935–41. Epub 2002/08/29. doi: 10.1038/nature00965. PubMed PMID: 12198538.

2. Roden LC, Carré IA. The molecular genetics of circadian rhythms in Arabidopsis. Semin Cell Dev Biol. 2001;12(4):305–15. doi: 10.1006/scdb.2001.0258. PubMed PMID: 11463215.

3. Xue Z, Ye Q, Anson SR, Yang J, Xiao G, Kowbel D, et al. Transcriptional interference by antisense RNA is required for circadian clock function. Nature. 2014;514(7524):650–3. Epub 2014/08/19. doi: 10.1038/nature13671. PubMed PMID: 25132551; PubMed Central PMCID: PMCPMC4214883.

4. Mori T, Johnson CH. Circadian programming in cyanobacteria. Semin Cell Dev Biol. 2001;12(4):271–8. Epub 2001/07/21. doi: 10.1006/scdb.2001.0254. PubMed PMID: 11463211.

5. Loros JJ, Dunlap JC. Genetic and molecular analysis of circadian rhythms in Neurospora. Annual review of physiology. 2001;63:757–94. Epub 2001/02/22. doi: 10.1146/annurev.physiol.63.1.757. PubMed PMID: 11181975.

6. Goldbeter A. Biochemical Oscillations and Cellular Rhythms: The Molecular Bases of Periodic and Chaotic Behaviour: Cambridge University Press; 1997.

7. Kurosawa G, Aihara K, Iwasa Y. A model for the circadian rhythm of cyanobacteria that maintains oscillation without gene expression. Biophys J. 2006;91(6):2015–23. Epub 2006/06/27. doi: 10.1529/biophysj.105.076554. PubMed PMID: 16798799; PubMed Central PMCID: PMCPMC1557573.

8. Goodwin BC. Oscillatory behavior in enzymatic control processes. Advances in enzyme regulation. 1965;3:425–38. Epub 1965/01/01. PubMed PMID: 5861813.

9. Gonze D, Abou-Jaoude W. The Goodwin model: behind the Hill function. PloS one. 2013;8(8):e69573. Epub 2013/08/13. doi: 10.1371/journal.pone.0069573. PubMed PMID: 23936338; PubMed Central PMCID: PMCPMC3731313.

10. Tyson JJ, Othmer HG. The dynamics of feedback control circuits in biochemical pathways. Prog Theor Biol. 1978;5:1–62.

11. Kim JK, Forger DB. A mechanism for robust circadian timekeeping via stoichiometric balance. Molecular systems biology. 2012;8:630. Epub 2012/12/06. doi: 10.1038/msb.2012.62. PubMed PMID: 23212247; PubMed Central PMCID: PMCPMC3542529.

12. Mirsky HP, Liu AC, Welsh DK, Kay SA, Doyle FJ. A model of the cell-autonomous mammalian circadian clock. Proceedings of the National Academy of Sciences. 2009;106(27):11107–12. doi: 10.1073/pnas.0904837106.

13. Leloup J-C, Goldbeter A. Toward a detailed computational model for the mammalian circadian clock. Proceedings of the National Academy of Sciences of the United States of America. 2003;100(12):7051–6. doi: 10.1073/pnas.1132112100. PubMed PMID: PMC165828.

14. Korenčič A, Bordyugov G, Košir R, Rozman D, Goličnik M, Herzel H. The Interplay of cis-Regulatory Elements Rules Circadian Rhythms in Mouse Liver. PloS one. 2012;7(11):e46835. doi: 10.1371/journal.pone.0046835. PubMed PMID: PMC3489864.

15. Hong CI, Jolma IW, Loros JJ, Dunlap JC, Ruoff P. Simulating Dark Expressions and Interactions of *frq* and *wc-1* in the *Neurospora* Circadian Clock. Biophysical Journal. 2008;94(4):1221–32. doi: 10.1529/biophysj.107.115154.

16. Relógio A, Westermark PO, Wallach T, Schellenberg K, Kramer A, Herzel H. Tuning the Mammalian Circadian Clock: Robust Synergy of Two Loops. PloS Computational Biology. 2011;7(12):e1002309. doi: 10.1371/journal.pcbi.1002309.

17. Yan J, Shi G, Zhang Z, Wu X, Liu Z, Xing L, et al. An intensity ratio of interlocking loops determines circadian period length. Nucleic Acids Research. 2014;42(16):10278–87. doi: 10.1093/nar/gku701.

18. Saithong T, Painter KJ, Millar AJ. The Contributions of Interlocking Loops and Extensive Nonlinearity to the Properties of Circadian Clock Models. PloS one. 2010;5(11):e13867. doi: 10.1371/journal.pone.0013867.

19. Pett JP, Korenčič A, Wesener F, Kramer A, Herzel H. Feedback Loops of the Mammalian Circadian Clock Constitute Repressilator. PloS Computational Biology. 2016;12(12):e1005266. doi: 10.1371/journal.pcbi.1005266. PubMed PMID: PMC5189953.

20. Foo M, Somers DE, Kim P-J. Kernel Architecture of the Genetic Circuitry of the Arabidopsis Circadian System. PloS Computational Biology. 2016;12(2):e1004748. doi: 10.1371/journal.pcbi.1004748.

21. Battogtokh D, Tyson JJ. Bifurcation analysis of a model of the budding yeast cell cycle. Chaos (Woodbury, NY). 2004;14(3):653–61. Epub 2004/09/28. doi: 10.1063/1.1780011. PubMed PMID: 15446975.

22. Anafi RC, Lee Y, Sato TK, Venkataraman A, Ramanathan C, Kavakli IH, et al. Machine Learning Helps Identify CHRONO as a Circadian Clock Component. PloS Biology. 2014;12(4):e1001840. doi: 10.1371/journal.pbio.1001840. PubMed PMID: PMC3988006.

23. Gérard C, Goldbeter A. Temporal self-organization of the cyclin/Cdk network driving the mammalian cell cycle. Proceedings of the National Academy of Sciences of the United States of America. 2009;106(51):21643–8. doi: 10.1073/pnas.0903827106. PubMed PMID: PMC2799800.

24. Battogtokh D, Aihara K, Tyson JJ. Synchronization of eukaryotic cells by periodic forcing. Physical review letters. 2006;96(14):148102. Epub 2006/05/23. doi: 10.1103/PhysRevLett.96.148102. PubMed PMID: 16712125.

25. Gerard C, Goldbeter A. Entrainment of the mammalian cell cycle by the circadian clock: modeling two coupled cellular rhythms. PloS Comput Biol. 2012;8(5):e1002516. Epub 2012/06/14. doi: 10.1371/journal.pcbi.1002516. PubMed PMID: 22693436; PubMed Central PMCID: PMCPMC3364934.

26. Hong CI, Zámborszky J, Baek M, Labiscsak L, Ju K, Lee H, et al. Circadian rhythms synchronize mitosis in Neurospora crassa. Proceedings of the National Academy of Sciences. 2014;111(4):1397–402. doi: 10.1073/pnas.1319399111.

27. Tyson JJ, Novák B. Functional Motifs in Biochemical Reaction Networks. Annual Review of Physical Chemistry. 2010;61(1):219–40. doi: 10.1146/annurev.physchem.012809.103457. PubMed PMID: 20055671.

28. Hartwell LH, Hopfield JJ, Leibler S, Murray AW. From molecular to modular cell biology. Nature. 1999.

29. Traynard P, Feillet C, Soliman S, Delaunay F, Fages F. Model-based investigation of the circadian clock and cell cycle coupling in mouse embryonic fibroblasts: Prediction of RevErb-alpha up-regulation during mitosis. Bio Systems. 2016;149:59–69. Epub 2016/07/23. doi: 10.1016/j.biosystems.2016.07.003. PubMed PMID: 27443484.

30. Bieler J, Cannavo R, Gustafson K, Gobet C, Gatfield D, Naef F. Robust synchronization of coupled circadian and cell cycle oscillators in single mammalian cells. Molecular systems biology. 2014;10:739. Epub 2014/07/17. doi: 10.15252/msb.20145218. PubMed PMID: 25028488; PubMed Central PMCID: PMCPMC4299496.

31. Kornmann B, Schaad O, Bujard H, Takahashi JS, Schibler U. System-Driven and Oscillator-Dependent Circadian Transcription in Mice with a Conditionally Active Liver Clock. PloS Biology. 2007;5(2):e34. doi: 10.1371/journal.pbio.0050034.

32. Battogtokh D, Kojima S, Tyson JJ. Modeling the interactions of sense and antisense Period transcripts in the mammalian circadian clock network. PloS Comput Biol. 2018;14(2):e1005957. Epub 2018/02/16. doi: 10.1371/journal.pcbi.1005957. PubMed PMID: 29447160.

33. Doedel E, Champneys A, Dercole F, Fairgrieve T, Kuznetsov Y, Oldeman B, et al. Auto: {S}oftware for continuation and bifurcation problems in ordinary differential equations. 2009.

34. Hughes ME, DiTacchio L, Hayes KR, Vollmers C, Pulivarthy S, Baggs JE, et al. Harmonics of Circadian Gene Transcription in Mammals. PloS genetics. 2009;5(4):e1000442. doi: 10.1371/journal.pgen.1000442. PubMed PMID: PMC2654964.

35. Relogio A, Thomas P, Medina-Perez P, Reischl S, Bervoets S, Gloc E, et al. Ras-mediated deregulation of the circadian clock in cancer. PloS genetics. 2014;10(5):e1004338. Epub 2014/05/31. doi: 10.1371/journal.pgen.1004338. PubMed PMID: 24875049; PubMed Central PMCID: PMCPMC4038477.

36. El-Athman R, Genov NN, Mazuch J, Zhang K, Yu Y, Fuhr L, et al. The Ink4a/Arf locus operates as a regulator of the circadian clock modulating RAS activity. PloS Biology. 2017;15(12):e2002940. doi: 10.1371/journal.pbio.2002940.

37. Dibner C, Sage D, Unser M, Bauer C, d’Eysmond T, Naef F, et al. Circadian gene expression is resilient to large fluctuations in overall transcription rates. The EMBO Journal. 2009;28(2):123–34. doi: 10.1038/emboj.2008.262. PubMed PMID: PMC2634731.

38. Gonze D, Bernard S, Waltermann C, Kramer A, Herzel H. Spontaneous Synchronization of Coupled Circadian Oscillators. Biophys J. 2005;89(1):120–9. doi: 10.1529/biophysj.104.058388. PubMed PMID: 15849258; PubMed Central PMCID: PMCPMC1366510.

39. Levi F, Schibler U. Circadian rhythms: mechanisms and therapeutic implications. Annu Rev Pharmacol Toxicol. 2007;47:593–628. Epub 2007/01/11. doi: 10.1146/annurev.pharmtox.47.120505.105208. PubMed PMID: 17209800.

40. Bugge A, Feng D, Everett LJ, Briggs ER, Mullican SE, Wang F, et al. Rev-erbalpha and Rev-erbbeta coordinately protect the circadian clock and normal metabolic function. Genes Dev. 2012;26(7):657–67. Epub 2012/04/05. doi: 10.1101/gad.186858.112. PubMed PMID: 22474260; PubMed Central PMCID: PMCPMC3323877.

41. Cho H, Zhao X, Hatori M, Yu RT, Barish GD, Lam MT, et al. Regulation of circadian behaviour and metabolism by REV-ERB-alpha and REV-ERB-beta. Nature. 2012;485(7396):123–7. Epub 2012/03/31. doi: 10.1038/nature11048. PubMed PMID: 22460952; PubMed Central PMCID: PMCPMC3367514.

42. Edwards MD, Brancaccio M, Chesham JE, Maywood ES, Hastings MH. Rhythmic expression of cryptochrome induces the circadian clock of arrhythmic suprachiasmatic nuclei through arginine vasopressin signaling. Proceedings of the National Academy of Sciences of the United States of America. 2016;113(10):2732–7. doi: 10.1073/pnas.1519044113. PubMed PMID: PMC4791030.

43. Johnson CH. Circadian clocks and cell division: What’s the pacemaker? Cell Cycle. 2010;9(19):3864–73. doi: 10.4161/cc.9.19.13205. PubMed PMID: PMC3047750.

44. Chao H-W, Doi M, Fustin J-M, Chen H, Murase K, Maeda Y, et al. Circadian clock regulates hepatic polyploidy by modulating Mkp1-Erk1/2 signaling pathway. Nature Communications. 2017;8:2238. doi: 10.1038/s41467-017-02207-7. PubMed PMID: PMC5740157.

45. Borisuk MT, Tyson JJ. Bifurcation analysis of a model of mitotic control in frog eggs. Journal of theoretical biology. 1998;195(1):69–85. Epub 1998/11/06. doi: 10.1006/jtbi.1998.0781. PubMed PMID: 9802951.

46. Lehmann R, Childs L, Thomas P, Abreu M, Fuhr L, Herzel H, et al. Assembly of a comprehensive regulatory network for the mammalian circadian clock: a bioinformatics approach. PloS one. 2015;10(5):e0126283. Epub 2015/05/07. doi: 10.1371/journal.pone.0126283. PubMed PMID: 25945798; PubMed Central PMCID: PMCPMC4422523.

